# The time course of different surround suppression mechanisms

**DOI:** 10.1101/432773

**Authors:** Michael-Paul Schallmo, Alex M. Kale, Scott O. Murray

**Affiliations:** Department of Psychology, University of Washington, Seattle, WA, USA; Department of Psychiatry and Behavioral Science, University of Minnesota, Minneapolis, MN, USA

## Abstract

What we see depends on the spatial context in which it appears. Previous work has linked the reduction of perceived stimulus contrast in the presence of surrounding stimuli to the suppression of neural responses in early visual cortex. It has also been suggested that this surround suppression depends on at least two separable neural mechanisms, one ‘low-level’ and one ‘higher-level,’ which can be differentiated by their response characteristics. In a recent study, we found evidence consistent with these two suppression mechanisms using psychophysical measurements of perceived contrast. Here, we used EEG to demonstrate for the first time that neural responses in the human occipital lobe also show evidence of two separable suppression mechanisms. Eighteen adults (10 female and 8 male) each participated in a total of 3 experimental sessions, in which they viewed visual stimuli through a mirror stereoscope. The first session was used to definitively identify the C1 component, while the second and third comprised the main experiment. ERPs were measured in response to center gratings either with no surround, or with surrounding gratings oriented parallel or orthogonal, and presented either in the same eye (monoptic) or opposite eye (dichoptic). We found that the earliest ERP component (C1; ∼60 ms) was suppressed in the presence of surrounding stimuli, but that this suppression did not depend on surround configuration, suggesting a low-level suppression mechanism which is not tuned for relative orientation. A later response component (N1; ∼160 ms) showed stronger surround suppression for parallel and monoptic stimulus configurations, consistent with our earlier psychophysical results and a higher-level, binocular, orientation-tuned suppression mechanism. We conclude that these two surround suppression mechanisms have distinct response time courses in the human visual system, which can be differentiated using electrophysiology.

## Introduction

The spatial context in which an image is seen dramatically affects both how it is perceived and the underlying neural response. However, the link between the perceptual effects of spatial context and the underlying neural mechanisms remains incomplete. Surround suppression is a spatial context phenomenon in which the presence of a surrounding stimulus reduces the neural response to a center stimulus, compared to when that center image is viewed in isolation. This effect is clearly observed when stimuli are presented both within and outside the classical receptive field of neurons in primary visual cortex (V1), as measured by electrophysiology in animal models (Bair, Cavanaugh, & Movshon, 2003; Cavanaugh, Bair, & Movshon, 2002a, 2002b; DeAngelis, Freeman, & Ohzawa, 1994; Ichida, Schwabe, Bressloff, & Angelucci, 2007; Shushruth, Ichida, Levitt, & Angelucci, 2009; Shushruth et al., 2013; Walker, Ohzawa, & Freeman, 1999). Surround suppression is also observed in human visual perception; the perceived contrast or discriminability of a center stimulus is reduced in the presence of a surround (Chubb, Sperling, & Solomon, 1989; Ejima & Takahashi, 1985; Petrov & McKee, 2006; Xing & Heeger, 2000, 2001; Yu, Klein, & Levi, 2001). Perceptual surround suppression is thought to depend on suppressed neural responses in human visual cortex (Schallmo, Grant, Burton, & Olman, 2016; Self et al., 2016; Zenger-Landolt & Heeger, 2003). Indeed, stimuli that produce perceptual surround suppression also evoke suppressed neural responses in the human occipital lobe, as measured by magneto-or electroencephalography (MEG or EEG; Applebaum, Wade, Vildavski, Pettet, & Norcia, 2006; Haynes, Roth, Stadler, & Heinze, 2003; Joo, Boynton, & Murray, 2012; Joo & Murray; Ohtani, Okamura, Yoshida, Toyama, & Ejima, 2002; Vanegas, Blangero, & Kelly, 2015). Typically, greater suppression is observed for more-similar center and surrounding stimuli (e.g., parallel orientation). It has been suggested that surround suppression may serve a number of different functional roles in visual processing, including supporting figure-ground segmentation (Poort et al., 2012; Poort, Self, van Vugt, Malkki, & Roelfsema, 2016; Roelfsema & de Lange, 2016), perceptual grouping (Joo et al., 2012; Joo & Murray, 2014), perceptual inference (Coen-Cagli, Kohn, & Schwartz, 2015), and efficient coding of information (Vinje & Gallant, 2000). Although much is known about the phenomenon of surround suppression, the neural mechanisms that give rise to this effect remain imperfectly understood.

The time course of surround suppression may provide insight into the underlying neural processes. The pioneering work of Bair and colleagues (2003) showed that surround suppression in macaque V1 has a latency of ∼61 ms, slightly longer than that for the onset of a response to the center stimulus (∼52 ms). Based on their results, they concluded that a fast feedback mechanism from higher visual areas (e.g., V2 or MT) may account for the observed time course of suppression. Using MEG, Ohtani and colleagues (2002) found that the earliest measured response to a center grating (∼90 ms) was suppressed (but not delayed) by the presence of a surrounding grating. Haynes and colleagues (2003) used both MEG and EEG to examine surround suppression, and compared neural response magnitudes to measures of perceived contrast. They found suppression for collinear vs. orthogonal center-surround configurations in both early (∼80 ms) and later (∼130 ms) response components, but that the later response was most closely associated with perception. From this early work and more recent studies (Chen, Yu, Zhu, Peng, & Fang, 2016; Joo et al., 2012; Joo & Murray; Miller, Shapiro, & Luck, 2015; Vanegas et al., 2015), it is clear that surround suppression can be observed in the earliest visually-evoked responses from occipital areas (e.g., V1). However, the precise neural origin of this suppression (i.e., feed-forward, later interactions, and / or feedback) is not yet clear.

It has been shown that surround suppression depends on at least two separable neural mechanisms with different physiological properties. Webb and colleagues (2005) used electrophysiology in macaque V1 to show that one form of suppression (hereafter referred to as ‘low-level’) is monocular, broadly tuned for orientation, and resistant to adaptation. They contrasted this with a second type of suppression (‘higher-level’ hereafter), which is binocular, stronger for parallel centers and surrounds (i.e., orientation selective), and greatly attenuated following 30 s of adaptation with a dynamic annular surround stimulus. This is consistent with current theories about the neural circuit origins of surround suppression in V1; some amount of non-selective suppression is thought to be inherited in a feed-forward manner from center-surround antagonism in the lateral geniculate nucleus (LGN), while horizontal connections within V1 and feedback from higher areas (e.g., V2, V4, MT) are thought to provide additional suppression that is selective for particular stimulus features (e.g., orientation; Angelucci & Bressloff, 2006; Nurminen & Angelucci, 2014). Psychophysical studies in humans suggest that perceptual surround suppression also depends on the same low-and higher-level mechanisms (Cai, Zhou, & Chen, 2008; Petrov & McKee, 2009). We recently showed that suppression of perceived contrast was stronger for parallel vs. orthogonal surrounds viewed dichoptically (i.e., through a stereoscope, with the center appearing in one eye and the surround in the other), consistent with a binocular, orientation-selective, higher-level suppression mechanism (Schallmo & Murray, 2016). We further showed that after adaptation in the surrounding region to a contrast-reversing grating, dichoptic suppression was eliminated, and monoptic suppression (i.e., within the same eye) was equivalent for parallel and orthogonal surrounds. This second result was consistent with a monocular, orientation non-selective, low-level form of suppression. Using the stereoscope permitted us to distinguish monocular vs. binocular processes, suggesting different stages of visual processing. Additional evidence of an inter-ocular “contrast normalization” mechanism that is orientation selective has been observed using functional MRI (Moradi & Heeger, 2009). However, to our knowledge, low- and higher-level suppression have not been observed in human visual cortex using electrophysiology. Thus, physiological evidence for these two suppression mechanisms in humans is lacking, and their particular neural time courses remain unknown.

The current study sought to fill this gap in our knowledge by quantifying the time signature of these two putative surround suppression mechanisms in the human visual system. We acquired event-related potential (ERP) measurements in a paradigm we developed previously (Schallmo & Murray, 2016) in an attempt to distinguish low- and higher-level processes (Angelucci & Bressloff, 2006; Nurminen & Angelucci, 2014; Webb et al., 2005). We expected that suppression by a low-level mechanism would be reflected in an ERP component occurring early in time, which would not be modulated by surround orientation, and that this suppression would be absent for dichoptic stimulus presentations. Further, we anticipated that higher-level suppression would manifest in a later ERP component, which would be sensitive to both surround orientation and eye-of-origin. Because the functional significance of particular ERP components is not fully understood, we did not have strong *a priori* expectations for how low- and higher-level suppression would map onto the visual components of interest (C1, P1, N1, and P2), beyond the notion that the C1 was the most likely to reflect low-level suppression, due to its short latency and putative locus in early visual cortex (i.e., V1; Clark, Fan, & Hillyard, 1994; Di Russo, Martinez, Sereno, Pitzalis, & Hillyard, 2002; Kelly, Schroeder, & Lalor, 2013); but see (Ales, Yates, & Norcia, 2010) for a dissenting view. Finally, we expected that higher-level suppression would manifest in a later component (e.g., N1 or P2).

## Methods

### Participants

Prior to data collection, we performed a power analysis to calculate the appropriate number of subjects for recruitment. Based on the smallest effect sizes and corresponding standard deviations from published EEG studies of surround suppression (Applebaum et al., 2006; Chicherov & Herzog; Chicherov, Plomp, & Herzog, 2014; Haynes et al., 2003; Joo et al., 2012; Joo & Murray; Silver, Kosovicheva, & Landau, 2011; Vanegas et al., 2015), we determined that 16 subjects would provide an *a priori* power of 92% for detecting a significant difference in surround suppression between experimental conditions. We initially assumed a conservative subject retention rate of 80%, and therefore recruited a total of 20 subjects (10 female and 10 male). Two of these subjects (both male) were ultimately excluded; one subject withdrew before completing the entire experiment, and another was excluded post hoc due to excessive alpha oscillations in all experimental sessions (based on visual inspection). Data from 18 subjects were included in the final analyses.

Subjects affirmed that they had normal or corrected-to-normal binocular vision. Individuals who exclusively wore contact lenses (and not eyeglasses) were screened out, as contacts are known to produce a high rate of eye blinks, which evoke EEG signal artifacts. Subjects provided written informed consent prior to participation and were compensated $20 per hour. All procedures were approved by the University of Washington Institutional Review Board (approval #35433) and complied with the Declaration of Helsinki.

### Apparatus

EEG data were collected using a Biosemi system with 64 active Ag-AgCl electrodes at a sampling rate of 256 Hz. Caps were positioned on each subject’s head with the Cz electrode placed half way between the nasion and the inion in the anterior-posterior dimension, and half way between the tragus of the left and right ears. During data collection, a Common Mode Sensor reference was used. Data were recorded on a PC running Windows XP using Biosemi’s ActiView software.

Visual stimuli were displayed on a ViewSonic P225f CRT monitor by a second PC running Windows XP using Presentation software (Neurobehavioral Systems, Berkeley, CA). Monitor luminance was linearized using a Photoresearch PR650 spectrophotometer and corrected within Presentation. A mirror stereoscope was used to present the left and right halves of the screen separately to the left and right eyes. The stereoscope reduced the maximum luminance of the moniator from 105 to 45 cd/m^2^. Prior to the start of each experimental session, the stereoscope was calibrated for each individual, and subjects performed a series of assessments to ensure stable fusion of the images presented to the two eyes, as in our previous study (Schallmo & Murray, 2016).

### Stimuli

Visual stimuli matched those used previously (Schallmo & Murray, 2016), except where noted below. Briefly, sinusoidal luminance modulation gratings were presented within circular apertures arranged into 3 x 3 square grids. The grating in the center of each grid constituted the target, while the gratings along the edge were referred to as surrounds. Target gratings were present in all experimental conditions, while the presence and configuration of surrounds varied between conditions (see below). The size, spatial frequency, and the position of the gratings differed slightly from our previous study (Schallmo & Murray, 2016), in order to facilitate detection of the C1 component (Clark et al., 1994; Di Russo et al., 2002). Gratings were presented within a circular aperture (radius 0.95°) blurred with a Gaussian envelope (SD = 0.05°). The center-to-center distance of adjacent gratings within the grid was 1.2°. All gratings were presented spatially in-phase with a spatial frequency of 4 cycles/°, oriented either vertically or horizontally.

There were 4 possible spatial locations on the screen where the stimulus grids could appear, two on the left half and two on the right. When viewed through the stereoscope, the left and right halves of the screen overlapped fully and merged into a single display, such that subjects perceived only two grids (Figure 1A). These grids were presented peripherally, centered at 4.96° to the left or right and 2° above a fixation mark at the middle of each screen-half (with one exception, see below). To aid binocular fusion, the fixation mark in each eye consisted of a high-contrast square pattern, intersected by a horizontal line in the left eye and a vertical line in the right eye (which formed a plus-sign when properly fused). Targets appeared within a thin fiducial circle.

**Figure 1.**
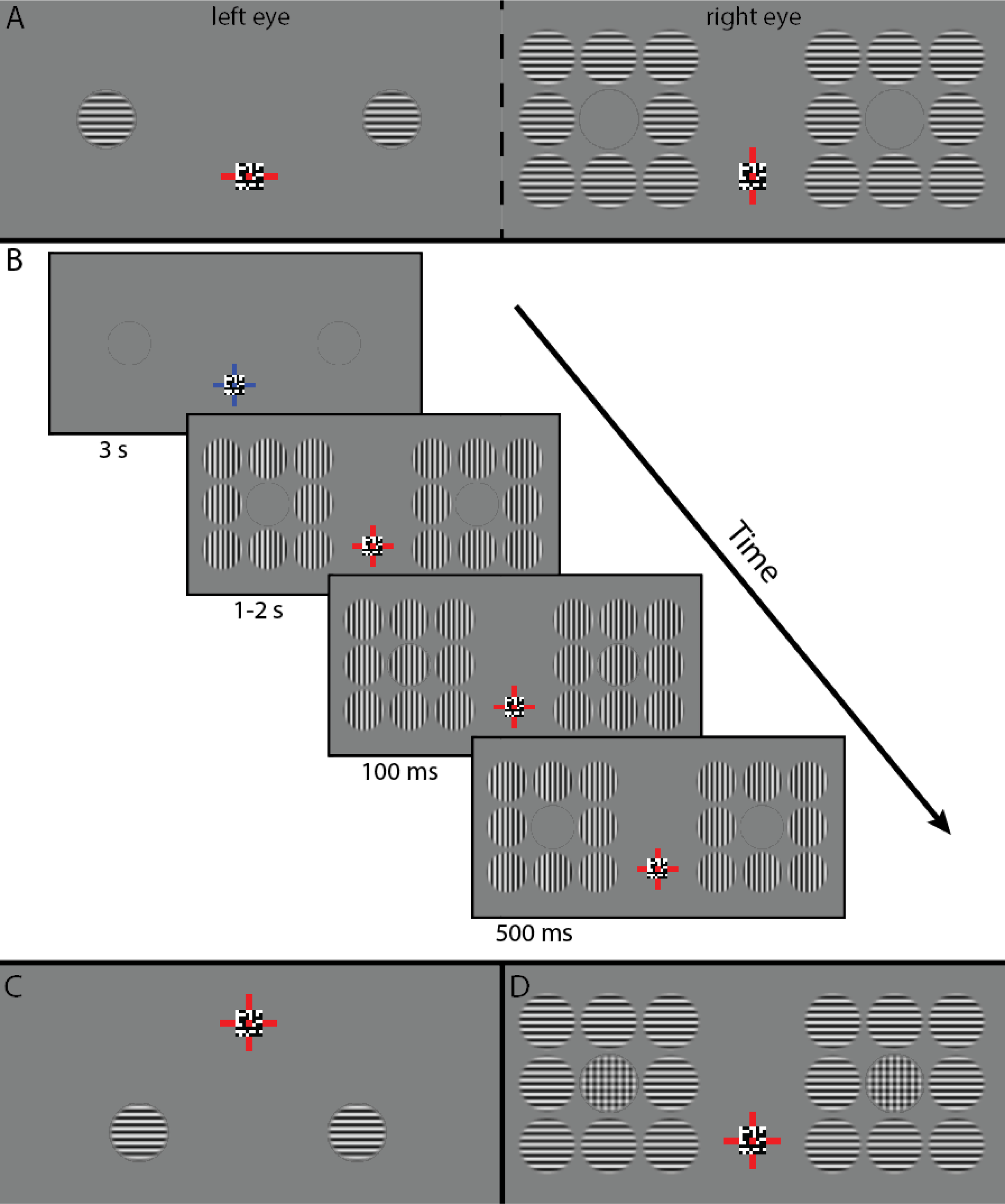
Stimuli and presentation timing. A. Dichoptic stimulus presentation. Left and right eyes see different images, when viewed through a mirror stereoscope, the two images fuse into a single percept. B. Stimulus presentation timing for a single trial. After a 3 s inter-trial interval, surrounds appear for 1-2 s, followed by the targets (center of 3 x 3 grid) for 100 ms. Surrounds remain in place for 500 ms after target offset before the end of the trial. C. Stimuli presented in the lower visual field as part of the C1 experiment. D. Plaid stimuli. The subjects’ task in the main experiment was to respond to the infrequent presentation of plaid stimuli (20% of trials).

### Paradigm

Subjects completed two separate experimental paradigms as part of this study: 1) the C1 experiment, designed to measure the earliest visual ERP component (known as C1), and 2) the main experiment, which characterized the ERP responses to different configurations of target and surround stimuli. Experimental methods common to both paradigms were as follows. Experiments consisted of a series of trials in which stimuli from a single experimental condition were presented (Figure 1B). Trials began with a change in the fixation mark (from blue to red), which informed the subject of the trial onset. This was followed by a variable delay (1-2 s, randomized), after which the target gratings were presented at the center of both stimulus grids (left and right of fixation). Target duration was 100 ms. Following target offset, there was a 500 ms time period in which the fixation mark remained red, prior to the end of the trial. Once the trial ended, the fixation mark changed back from red to blue, and there was a delay period (inter-trial interval) of 3 s before the next trial began. Subjects were instructed to withhold blinks until the inter-trial interval whenever possible. 50 trials comprised one block, with a duration of about 4 min. Subjects received a short, self-timed break between blocks in order to rest their eyes. A single experimental session included 8-13 blocks (see below), and lasted approximately 1 hr. Further details specific to each experiment are provided below.

#### C1 experiment

This experiment was designed to identify the time window and scalp topography of the earliest visual ERP component, known as the C1, within our group of subjects. This experiment consisted of two conditions: 1) upper visual field, in which targets were presented at the aforementioned position (4.96° away horizontally, and 2° above the fixation mark), and 2) lower visual field, in which targets appeared below fixation (offset 3.78° horizontally and vertically; Figure 1C). These positions were chosen to try and maximize the difference in C1 polarity observed for stimuli presented in the upper (negative) and lower (positive) visual fields (Clark et al., 1994; Di Russo et al., 2002). Upper and lower visual field trials were presented in separate experimental blocks within the same session, to ensure that subjects knew where on the screen the targets would appear on every trial. Stimuli were presented binocularly in this experiment (i.e., shown on both the left and right halves of the screen, and seen in both the left and right eye).

In both the upper and lower visual field conditions, target gratings were oriented either vertically or horizontally; subjects were instructed to attend to the target orientation and report on each trial whether the target that appeared matched the orientation of the target on the previous trial or not (1-back task). Responses were made using mouse buttons, and subjects were instructed to wait until after each trial ended to respond. Twenty-five trials of each orientation were presented in each block in a random order, and subjects completed a total of 8 blocks within a single session for the C1 experiment (4 upper and 4 lower visual field, 200 trials total in each of these conditions). The order of upper vs. lower visual field blocks (first 4 or last 4) was counterbalanced across subjects.

#### Main experiment

This experiment was designed to measure the effect of surrounding stimuli on the ERP response to the target gratings. As in our previous study (Schallmo & Murray, 2016), surround orientation and optical configuration were manipulated in order to examine two putative neural mechanisms that contribute to surround suppression (Nurminen & Angelucci, 2014; Webb et al., 2005): a low-level, monoptic, and broadly orientation-tuned process, and a higher-level, binocular, narrowly orientation selective process. To this end, there were 6 different experimental conditions in the current study with different stimulus configurations: 1) no surround (target only), 2) parallel monoptic surround, 3) orthogonal monoptic surround, 4) parallel dichoptic surround, 5) orthogonal dichoptic surround, and 6) plaid target (part of the subjects’ task and used in a control analysis, see below). Parallel and orthogonal refer to the relative orientation between target and surrounds. Monoptic indicates that target and surrounds were presented to the same eye (i.e., on the same side of the monitor), while dichoptic indicates that target and surround were presented to opposite eyes.

For conditions with surrounding stimuli in the main experiments, surrounds were presented continuously throughout the trial (1-2 s before to 500 ms after the target presentation). This was done in order to isolate the ERP response to the target from any response to the surrounds, as in previous studies (Haynes et al., 2003; Joo et al., 2012; Joo & Murray, 2014; Ohtani et al., 2002). Stimuli in the main experiment were always presented in the upper visual field. The target orientation (vertical or horizontal) was randomized across trials and counterbalanced across conditions, as was the eye in which the target was presented (i.e., left or right side of the screen). There were 8 trials per condition in each block, except for the plaid condition (10 trials). All subjects completed 13 blocks in each session of the main experiment, except one subject who completed a 14^th^ block in one session, due to problems with signal artifacts during an earlier block. Each subject completed 2 sessions of the main experiment on separate days (the C1 session was always conducted on a another separate day, prior to the main experiment). Across both sessions, subjects completed a total of 208 trials in each of the main experiment conditions, except for the plaid condition (260 trials).

The subjects’ task in the main experiment was to respond to the appearance of plaid target stimuli (combined vertical and horizontal gratings; Figure 1D). Responses were made with a mouse button press after the end of the trial. This task served two purposes. First, it ensured that subjects attended to the target as it was presented. Second, the plaid condition served as a positive control; because plaid stimuli were presented in only 20% of the trials, we expected to see a strong P300 ERP response in this condition (Polich, 2007). While not the primary goal of our experiment, observing the expected P300 served to increase our confidence in the validity of the experimental procedure. In 80% of the plaid trials, surrounding stimuli were also presented as above (vertical or horizontal, monoptic or dichoptic with respect to the target, order randomized and counterbalanced across trials). Therefore, the presence, absence, or configuration of the surround did not serve as a cue for whether or not a plaid target would appear.

### Data processing and analysis

Data were processed and analyzed using an in-house toolbox written in MATLAB (The Mathworks, Natick, MA). Signals were re-referenced to the average of all electrodes. Trials were examined within epochs from -250 to +550 ms relative to target stimulus onset. High-pass filtering was not performed, as we found during data analysis that this introduced artifacts due to the onset and offset of the surrounding stimuli, which preceded and followed the target in time. Low-pass filtering was performed at 50 Hz. To exclude ringing artifacts from the low-pass filter, the first and last 50 ms of each epoch were not analyzed further. Hence, the epoch for data analysis and visualization was -200 to +500 ms. Signals within each epoch were baseline corrected by subtracting the mean signal between -200 and 0 ms (baseline window defined *a priori*) from all time points.

We were not able to collect EOG data for eyeblink artifact screening, due to the physical constraints of the stereoscope. Thus, eye movement and blink artifacts were detected using the following criteria defined *a priori*: epochs containing signal deflections > 75 µV or < -75 µV, relative to baseline, within electrodes F7, F8, and FPz (located at the anterior edge of the cap). The same criteria were used to define artifacts on electrodes Oz, O1, O2, POz, PO3, and PO4, which would be expected to result from head motion. Epochs with such artifacts were excluded; on average, 4.5% of trials were excluded in this way for the C1 experiment (range 0.3 - 19.5% across subjects), while 3.7% of trials were excluded in the main experiment (range 0 - 18.6%).

Four ERP components were selected *a priori* for analyses: C1, P1, N1, and P2. Electrode sites for ERP analyses were chosen *a priori* (Clark et al., 1994; Joo et al., 2012; Joo & Murray, 2014), and confirmed by visual inspection of scalp topographies (Supplemental Figure 1). For the C1, N1, and P2 components, 6 electrodes of interest were chosen along the midline (Oz, O1, O2, POz, PO3, and PO4). For the P1 component, which has a more lateral scalp distribution, electrodes P5, P6, P7, P8, PO7, and PO8 were chosen (Clark et al., 1994). EEG signals were averaged across these electrodes (separately for each time point within the components (in ms). These time windows epoch, each experimental condition, and each experimental session).

Time windows for each component were defined quantitatively based on data from the C1 experiment, within broad initial time windows defined *a priori*. These time windows were used for analysis of both the C1 experiment and the main experiment data. Consistent ERP timing was expected between the C1 experiment and the main experiment, as the spatial and temporal parameters of the stimuli were consistent between them. Thus, the C1 experiment data served as an independent localizer for component timing in the main experiment. The time windows for each component are shown in Table 1. The details of the analysis used in defining these time windows is provided below.

**Table 1.** Time windows for ERP components (in ms). These time windows were defined quantitatively from data in the C1 experiment.

|  | C1 | P1 | N1 | P2 |
| --- | --- | --- | --- | --- |
| **Start** | 61.5 | 88.8 | 159.1 | 217.7 |
| **End** | 84.9 | 135.7 | 206.0 | 256.8 |

To define ERP component time windows, we computed average time courses (across trials) for both the upper and lower visual field conditions in each subject, and took the integral of the EEG signal between each time point. For the C1 component, we calculated the response difference between the upper and lower visual field conditions (integrated EEG signal amplitude) at each time point, within each subject. This was done because the polarity of the C1 is known to depend on the position within the visual field in which the stimuli appear (upper = negative, lower = positive, as observed; Figure 2A) (Clark et al., 1994; Di Russo et al., 2002). We then averaged the integrated signal difference at each time point across subjects and found the maximum value (across time) of this average within the broad initial time window. The C1 window was defined by the time points at which the group average C1 difference (lower minus upper) was > 30% of the maximum. For the P1, N1, and P2 components, a similar procedure was used, but only the data from the upper visual field condition was used, and later initial time windows were applied.

**Figure 2.**
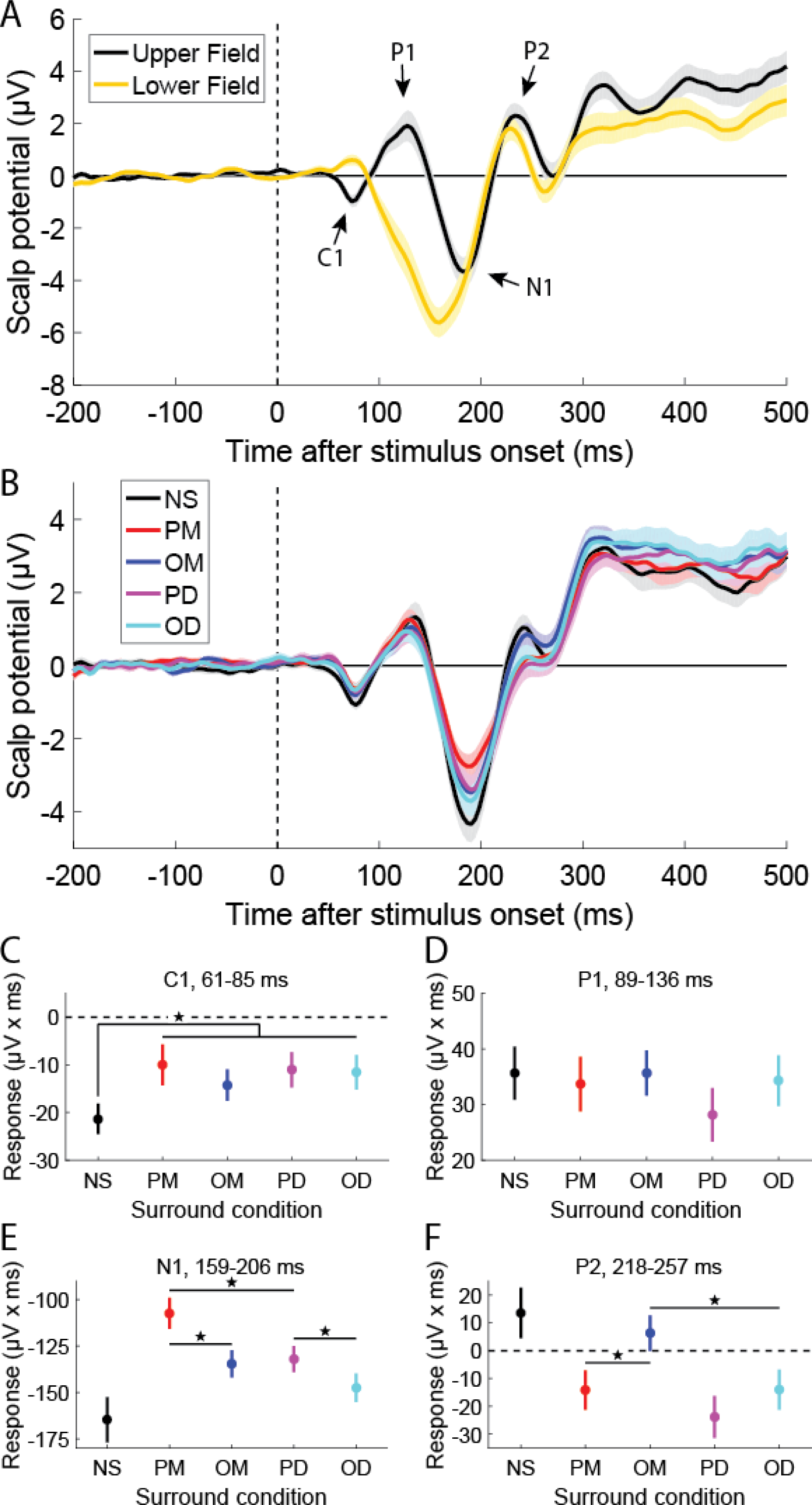
ERP results. **A**. Results from the C1 experiment; group-average ERPs for upper (black) vs. lower (yellow) visual field. Note the polarity inversion of the C1 between the two conditions. **B**. ERPs from the main experiment (NS: no surround, black; PM parallel monoptic, red; OM orthogonal monoptic, blue; PD parallel dichoptic, magenta; OD orthogonal dichoptic, cyan). ERP responses (integral of scalp potential) in each condition are shown for the C1 (**C**), P1 (**D**), N1 (**E**), and P2 (**F**). Error bars in **A** & **B** are *S.E.M.*, in **C-F**, within-subject error bars (Morey, 2008) are shown, to clarify differences between conditions. Asterisks indicate significant post-hoc paired *t*-tests, *p* < 0.05. Note the y-axes differ in **C-F**.

We did not expect any substantial differences between ERPs for vertical and horizontal target gratings, and no substantial differences were noted during our initial analyses, thus we averaged across target orientations for all subsequent analyses. Likewise, we did not expect or observe substantial differences between ERPs for stimuli presented on the left versus right side of the screen (when averaging across bilateral electrodes), and thus we subsequently averaged across screen side as well.

### Statistics

Statistical analyses were performed in MATLAB. The integrals of the scalp potentials measured within the time windows in Table 1 were used for all analyses. We performed mixed repeated measures ANOVAs (2 sessions per subject) for each ERP component. In first-level ANOVAs, the integrated scalp potential for each of the 5 stimulus conditions was included. For those ERP components in which a significant main effect of condition was observed, we performed a series of planned post-hoc tests to assess differences between particular surround conditions. Here, we compared the integrated signal magnitude in each condition using paired 2-tailed t-tests. We also performed an additional ANOVA to compare the effect of surround orientation (parallel vs. orthogonal) and optical configuration (monoptic vs. dichoptic); the no surround condition was not included in the post-hoc ANOVA. We sought to address the issue of multiple comparisons by performing post-hoc tests only in cases where a significant main effect of surround condition was observed in the first-level ANOVA.

## Results

We performed two experiments in a group of 18 subjects to characterize the ERP responses to different configurations of visual stimuli. The C1 experiment was designed to definitively identify the C1 component within our paradigm by measuring a polarity inversion for target stimuli presented in the upper vs. lower visual field (Clark et al., 1994; Di Russo et al., 2002). This experiment also served as an independent data set in which to define ERP component timing for our main experiment. The purpose of the main experiment was to measure the effect of surrounding stimuli on ERP responses to target gratings that were centered within a 3 x 3 grid (Figure 1B). In this experiment, surrounding stimuli appeared first in time, followed by briefly presented targets. This allowed us to test the hypothesis that separable low- and high-level neural mechanisms with different functional properties contribute to the phenomenon of surround suppression (Nurminen & Angelucci, 2014; Schallmo & Murray, 2016; Webb et al., 2005).

### C1 experiment

In the C1 experiment, we presented gratings in both the upper and lower visual field, in separate blocks of trials. We observed the expected polarity inversion for an early ERP component (Figure 2A); scalp potentials over middle occipital cortex were negative for stimuli in the upper visual field, and positive for those in the lower visual field. This difference was statistically significant (F1,17 = 24.2, *p* = 1 x 10^-4^), and consistent with the known visual field selectivity of the C1 component, which is thought to originate from retinotopic early visual cortex (e.g., V1; Clark et al., 1994; Di Russo et al., 2002).

We also identified 3 additional ERP components of interest, based on the combined inspection of scalp topographies (Supplemental Figure 1) and ERP time courses in the upper visual field condition (the location where stimuli were presented in the main experiment). Specifically, we saw a lateral occipital P1, a middle occipital N1, and a later middle occipital P2. The time windows for these ERP components (Table 1) identified in the C1 experiment were also used to analyze data from the main experiment. In this way, the C1 experiment served as an independent temporal localizer for ERP components, which allowed us to avoid circular definitions of ERP time windows within the main experiment dataset.

Subjects performed a 1-back task during this experiment and reported whether the orientation of the target (vertical or horizontal) matched that on the previous trial. Accuracy on this task was very high for all subjects (mean = 98.8%, *SD* = 2.6%), suggesting they were attending to the target stimuli as instructed.

### Main experiment

In the main experiment, we expected that the presence of surrounding gratings would attenuate ERP responses to the presentation of the target stimuli (i.e., surround suppression; Applebaum et al., 2006; Haynes et al., 2003; Joo et al., 2012; Joo & Murray; Ohtani et al., 2002; Vanegas et al., 2015). More specifically, we expected 2 primary results: first, that the earliest ERP component (C1) would be attenuated in the presence of monoptic but not dichoptic surrounds, and would not differ with surround orientation. Such a pattern would be consistent with a low-level surround suppression mechanism that is monocular, not strongly orientation selective, and occurs early in time (Angelucci & Bressloff, 2006; Nurminen & Angelucci, 2014; Webb et al., 2005). Second, we expected that a later ERP component (P1, N1, or P2) would be suppressed more strongly for both monoptic and parallel surround configurations (vs. dichoptic and orthogonal, respectively), consistent with a higher-level mechanism that is orientation selective and occurs later in time.

For the C1 component, we found that ERPs were significantly suppressed (less negative) for all surround conditions vs. no surround (main effect of condition, F4,68 = 2.79, *p* = 0.033; post-hoc tests, t35 > 2.38, *p* < 0.023; Figure 2C). However, we observed no significant differences between different surround conditions, including no difference between monoptic and dichoptic surrounds (main effect of optical configuration, F117 = 0.09, *p* = 0.8; post-hoc t35 > 0.74, *p* > 0.4). This pattern is only a partial match to the first expected result listed above, suggesting the time course of low-level suppression is more complex than predicted.

P1 signals did not vary significantly across surround conditions (main effect, F4,68 = 0.87, *p* = 0.5; Figure 2D), and thus were not analyzed further.

In contrast, the size of the N1 signal depended significantly on the surround condition; the N1 was suppressed for all conditions vs. no surround (main effect of condition, F4,68 = 7.84, *p* = 3 x 10^-5^; post-hoc t35 > 2.98, *p* < 0.005), with the exception of the orthogonal dichoptic condition (t35 = 1.58, *p* = 0.12). Further, N1 ERPs were more strongly suppressed for both monoptic and parallel conditions vs. dichoptic and orthogonal, respectively (main effect of orientation, F117 = 8.71, *p* = 0.009; post-hoc t35 > 2.19, *p* < 0.035; main effect of optical configuration, F117 = 12.3, *p* = 0.003; post-hoc t35 = 1.95, *p* < 0.060; Figure 2E). This pattern fits our second expected result listed above, and closely matches the pattern of perceived contrast suppression we observed previously in a comparable psychophysical paradigm (Schallmo & Murray, 2016).

For the P2, we again saw that ERP signal amplitudes varied significantly across conditions (main effect, F4,17 = 5.84, *p* = 4 x 10^-4^; Figure 2F); compared with the no surround condition, signals were suppressed in all other conditions (post-hoc tests, t35 > 3.28, *p* < 0.002) aside from the orthogonal monoptic (t35 = 0.95, *p* = 0.3). Suppression was again stronger for parallel vs. orthogonal configurations in the P2 (F117 = 6.12, *p* = 0.024), as expected. However, we found that for the P2 suppression was actually *stronger* for the dichoptic vs. monoptic conditions (main effect of optical configuration, F117 = 5.84, *p* = 0.027). We did not expect this particular effect but offer a possible interpretation in the Discussion.

The subjects’ task in the main experiment was to detect the presentation of a plaid target grating (which occurred in 20% of trials). Task performance was once again near perfect (mean hit rate = 97.5%, *SD* = 2.2%; mean correct rejection rate = 98.0%, *SD* = 2.1%), indicating that subjects were attending to the stimuli being presented in the target location, and could perceive them accurately. We also calculated ERP responses to the plaid targets, and observed a large P300 response (Supplemental Figure 2). This was expected, given that the task-relevant plaid stimuli were presented infrequently (20% of trials; Polich, 2007). These observations suggest that subjects were actively performing the task and were attending to the target stimuli as instructed.

## Discussion

We used ERPs to examine the time course of surround suppression during early visual processing. It is thought that this form of contextual modulation depends on more than one neural process, based on work in both animal models and humans (Angelucci & Bressloff, 2006; Cai et al., 2008; Nurminen & Angelucci, 2014; Petrov & McKee, 2009; Webb et al., 2005). Thus, we sought to determine whether two distinct neural mechanisms (Schallmo & Murray, 2016), one ‘low-level’ and another ‘higher-level,’ might be reflected in different patterns of ERP suppression across time for particular configurations of center and surrounding stimuli. We observed surround suppression of the earliest ERP component (C1); suppression of the EEG signal was reliably detected by about 60 ms post-stimulus, indicating that surrounding stimuli attenuate the earliest neural responses to visual stimuli that may be measured with this technique. After about 160 ms post-stimulus, we found significant modulation of the N1 component by different configurations of surrounding stimuli. In particular, suppression was orientation-tuned (stronger for parallel vs. orthogonal surrounds), and stronger within-vs. between-eyes. The pattern of surround suppression we observed in the N1 signal corresponds very closely to our previous measurements of perceptual suppression using the same stimulus configurations (Schallmo & Murray, 2016). Together, these two observations indicate that broadly-tuned surround suppression occurs early in time, with stimulus-selective suppression emerging later.

The observed time course of ERP suppression provides us insight into the neural mechanisms underlying this spatial context phenomenon. Based on the hypothesized low-level and higher-level mechanisms noted above, we predicted that we would observe two distinct patterns of suppression. For the earliest responses (C1), we considered two competing hypotheses. For the first, we predicted that we would observe no difference between parallel and orthogonal surround conditions (i.e., orientation insensitivity), whereas we expected that suppression would be stronger within-versus between-eyes. This prediction reflects the type of ‘low-level’ surround suppression that has been described in the literature as occurring early in time; it is both insensitive to the relative orientation of center and surround and monocular (Webb et al., 2005). This would be consistent with feed-forward suppression from the LGN to the input layers of V1 (Angelucci & Bressloff, 2006; Nurminen & Angelucci, 2014), as well as the pattern of surround suppression we observed psychophysically following adaptation of the surround region in our previous work (Schallmo & Murray, 2016). Alternatively, we considered whether C1 suppression might indeed show selectivity for surround orientation and eye-of-origin; this would be consistent with findings in macaque V1 showing fast, orientation selective surround suppression, possibly driven by feedback acting at the earliest stage of cortical visual processing (Bair et al., 2003). We would expect this second prediction to hold if ‘low-level’ suppression occurs only in the LGN, and ‘higher-level’ suppression is in-place by the time the C1 response is initiated.

Our current results agreed with our first prediction above, but only in part (Figure 2C); we observed that surround suppression in the C1 signal did not depend on surrounding stimulus orientation, but we did not find any difference in C1 amplitude between monoptic and dichoptic conditions. Thus, surround suppression of the C1 was equally strong within-vs. between-eyes. This suggests 3 possible interpretations. First, by the time of the C1 (60 ms after target onset), an early surround suppression mechanism may operate binocularly, but does not yet show significant orientation tuning. Second, dichoptic surround suppression in the C1 may reflect a binocular feedback mechanism (Angelucci & Bressloff, 2006; Chen et al., 2016; Nurminen & Angelucci, 2014), which is not orientation selective. Finally, C1 surround suppression may indeed be stronger for monoptic vs. dichoptic condition, but we do not have enough statistical power to detect this difference. The second and third notions appear less plausible. It has been suggested that C1 suppression may be affected by feedback modulation (e.g., by effortful attention; Chen et al., 2016), but see (Rauss, Schwartz, & Pourtois, 2011). Since the surrounding gratings appeared 1-2 s before the target, it is reasonable to consider whether a feedback mechanism may have already been engaged by the time the target appeared. However, it has been shown that feedback projections (e.g., from V2 to V1) which contribute to surround suppression (Nassi, Gomez-Laberge, Kreiman, & Born, 2014; Nassi, Lomber, & Born, 2013; Nurminen, Merlin, Bijanzadeh, Federer, & Angelucci, 2018) are orientation specific, with a patchy distribution that depends on preferred orientation (Angelucci & Bressloff, 2006; Marques, Nguyen, Fioreze, & Petreanu, 2018). It is thought that this orientation-tuned feedback drives stronger suppression for iso-oriented surrounding stimuli (Angelucci & Bressloff, 2006; Nurminen & Angelucci, 2014; Webb et al., 2005). Therefore, we would expect that if feedback was responsible for dichoptic C1 suppression, this suppression would be stronger for parallel vs. orthogonal surrounds. With regard to statistical power, both our *a priori* power analysis (see Methods - Participants), and the observed difference in C1 amplitudes between no surround and all with-surround conditions suggest that we should have had sufficient power to observe a difference in C1 suppression between monoptic and dichoptic surround conditions if one existed. Thus, although we did not expect that surround suppression of the C1 response would be insensitive to both relative orientation and eye-of-origin, we speculate that this reflects an intermediate-early stage of surround modulation for which suppression operates between eyes but is not yet orientation selective.

In the current study, we took care to definitively identify the C1 by comparing ERPs to stimuli in the upper vs. lower visual field in our first (C1) experiment (Figure 2A; Supplemental Figure 1B &C). The observed ERP polarity inversion between these conditions at ∼60 ms reflects the retinotopic selectivity which is the signature of the C1 response. While there is some controversy regarding whether such inversion indicates an origin for the C1 within area V1 *per se*, versus other extrastriate areas (e.g., V2; Ales et al., 2010; Clark et al., 1994; Di Russo et al., 2002; Kelly et al., 2013), we can reasonably assert that the measured C1 responses reflect a very early stage of visual processing within a medial region of retinotopic visual cortex.

We additionally hypothesized that surround suppression in later ERP components (e.g., N1) would show both greater suppression for parallel vs. orthogonal and monoptic vs. dichoptic surrounds, in line with our previous psychophysical results (Schallmo & Murray, 2016), and with a ‘higher-level’ suppression mechanism that is selective for both orientation and eye-of-origin. Our N1 results aligned very closely with this prediction (Figure 2E), suggesting that ‘higher-level’ suppression is reflected first in time in the N1 as measured by EEG. The timing (∼160 ms) and topography (Supplemental Figure 1E) of this N1 signal both point to a locus for N1 surround suppression within ventrolateral early visual cortex, possibly in extrastriate area(s) such as V4 (Di Russo et al., 2002). This notion is consistent with the idea that feature-selective ‘higher-level’ surround suppression operates via feedback from higher to lower visual areas (Nurminen & Angelucci, 2014; Webb et al., 2005). The orientation-selective surround suppression we observed in the N1 response is also generally consistent with the timing of figure-ground modulation observed in neural responses within primate visual cortex (e.g., V1 & V4) measured electrophysiologically (Poort et al., 2016; Roelfsema & de Lange, 2016). Because the pattern of surround suppression we observed in the N1 response across stimulus conditions so closely matched the psychophysical results from our earlier study (compare Figure 2E in the current study to Figure 2A from (Schallmo & Murray, 2016)), we speculate that the neural responses involved in generating the N1 signal may play an important role in the perceptual judgment of center contrast within this paradigm. This notion is consistent with previous work showing that N1 signals are associated with visual discrimination (Vogel & Luck, 2000).

In the current study, P1 responses to targets (∼90 ms) were not modulated by the presence or the configuration of the surrounding stimuli. Previous work from our group has shown that P1 responses are significantly modulated by different configurations of surrounding stimuli (e.g., parallel vs. orthogonal), consistent with a grouping processes in which suppression is stronger for stimuli that are grouped together perceptually (Joo et al., 2012; Joo & Murray, 2014). There are a number of methodological issues that can be ruled out. The lack of P1 modulation by surrounding stimuli in the current results is not a side effect of choosing lateral occipital / parietal electrodes; equivalent P1 results were obtained using mid-occipital electrode sites (as was done in the previous studies; not shown). Further, we used a very similar stimulus presentation paradigm (Figure 1B) and data acquisition protocol, with the addition of the stereoscope. The most plausible explanation seems to be stimulus differences; the current stimuli were somewhat larger (circular aperture with radius = 0.95° vs. Gaussian with σ = 0.72°), higher in spatial frequency (4 vs. 2 cycles/°), and targets were centered within a 3 x 3 grid of surrounding gratings (rather than within a linear array of 5). It seems plausible that some or all of these stimulus differences might have contributed to the size and sensitivity to surround modulation of the P1 response. Indeed, we observed that in our current C1 experiment, stimuli in the lower visual field produced little or no discernable P1 (Figure 2A), indicating that this component was sensitive to particular stimulus configurations in our paradigm.

We did not have strong *a priori* expectations about the pattern of the P2 response across stimulus conditions, apart from expecting significant surround suppression that was modulated by relative orientation and optical configuration in a manner similar to the N1. In general, this is what we found (Figure 2F), surrounding stimuli suppressed the P2 response, and this was stronger for parallel vs. orthogonal surrounds. One unexpected finding was that P2 suppression was actually stronger for dichoptic vs. monoptic surrounds, which is the inverse of what we found for the N1. Also unlike the N1, the P2 pattern did not match the psychophysical results from our earlier study (Schallmo & Murray, 2016). We speculate that stronger dichoptic P2 suppression may reflect a later stage process for resolving perceptual ambiguity when conflicting images are presented in each eye (i.e., target grating and one, but a blank gray center region in the other). This may be related to the process(es) that give rise to binocular rivalry (although our stimuli did not evoke significant rivalry themselves). Consistent with this idea, Mishra and colleagues (2009) found larger P2 signal modulation for dichoptic vs. monoptic viewing conditions in a study of binocular rivalry and attention. They speculated that this might reflect greater involvement of higher cortical areas in attentional processing to resolve stimulus conflict during dichoptic viewing. Future experiments may seek to more precisely characterize the role of P2 signals in resolving inter-ocular conflict by parametrically varying the congruence of stimuli presented dichoptically.

## Acknowledgements

We thank Mark Pettet for technical assistance in developing the EEG protocol and data analysis. We thank Anastasia V. Flevaris for helpful discussions regarding experiment design.

This work was supported by the National Institute of Health (F32 EY025121 to MPS, R01 MH106520 to SOM, T32 EY007031), and the University of Washington (Electrophysiology Research Fund to SOM).

## Supplemental information

**Supplemental Figure 1.**
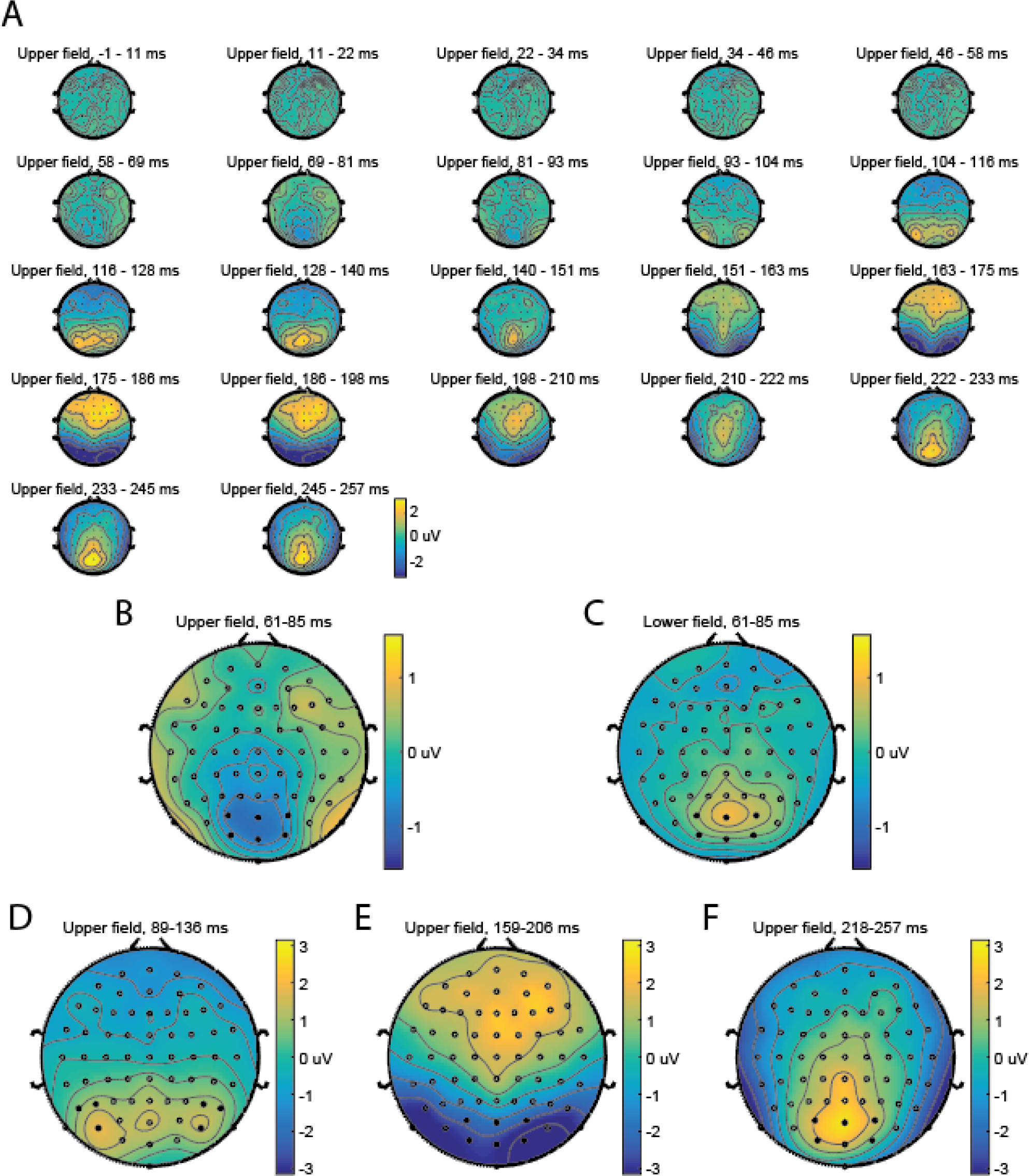
Scalp topographies. **A** shows scalp topographies for stimuli in the upper visual field from the C1 experiment across 22 evenly spaced time bins (with 3 EEG measurements per bin, averaged across N = 18 subjects). Topography plots were generated using the *topoplot.m* function from the EEGLAB toolbox for MATLAB (sccn.ucsd.edu/eeglab/). **B** & **C** show the average topographies for stimuli in the upper (**B**) and lower (**C**) visual field during the C1 time window. Note the polarity inversion of the signal near the occipital pole. Electrodes selected for C1 signal analyses (Oz, O1, O2, POz, PO3, PO4) are filled in black, while the rest are shown as open circles. **D-F** show the average topographies for the P1, N1, and P2 time windows, respectively. Note that the color scale differs from **B** & **C**. For the P1 window shown in **D**, a different set of electrodes were chosen (P5, P6, P7, P8, PO7, and PO8), and are filled in black.

**Supplemental Figure 2.**
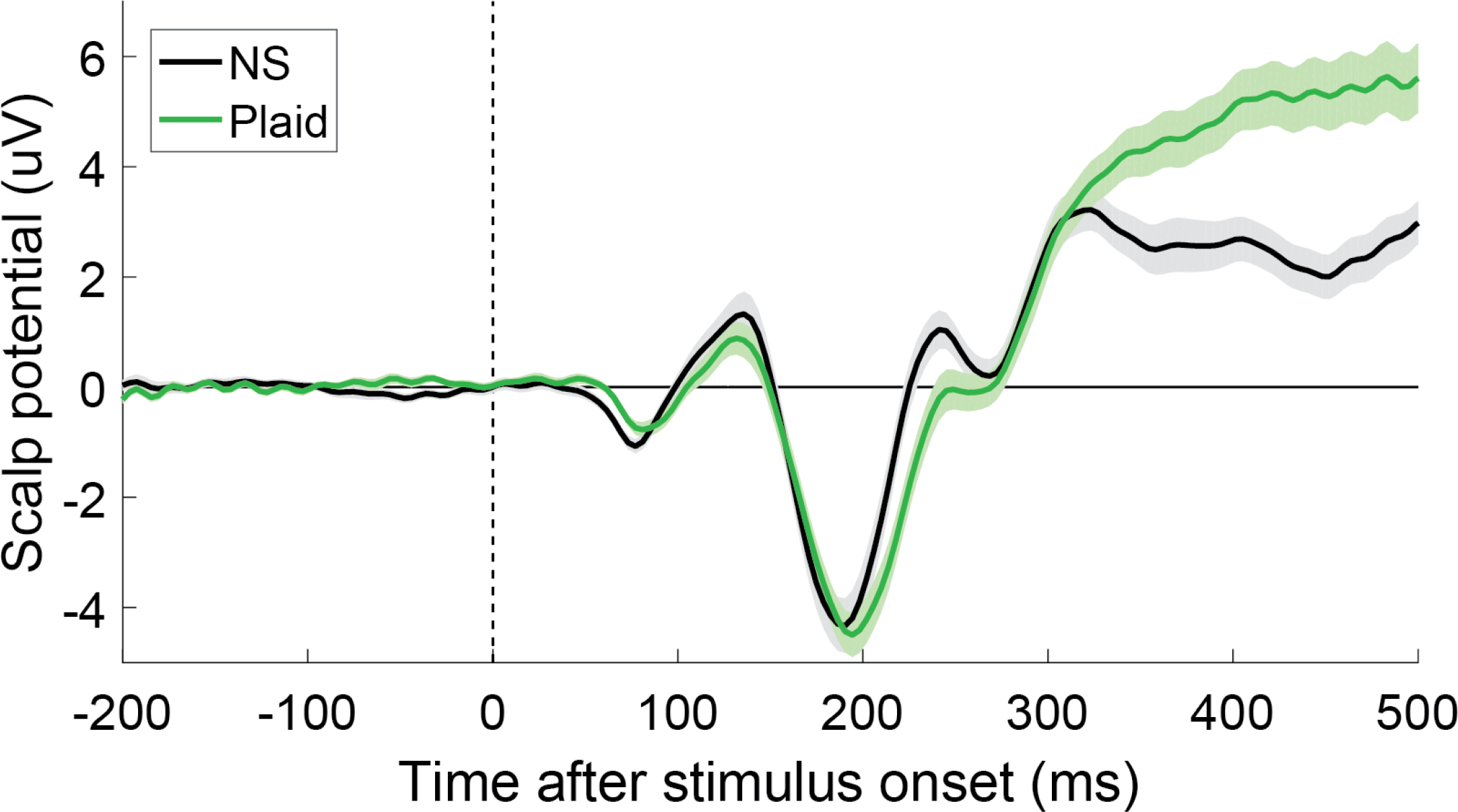
ERP response to the plaid stimuli. A P300 signal (positive deflection starting around 300 ms) is clearly visible in the subject-average ERP for stimuli in the plaid condition (green), as compared to those in the no surround (NS; black). These ERPs were recorded from occipital electrode sites (Oz, O1, O2, POz, PO3, PO4); a similar P300 was also observed over central-parietal electrodes (e.g., CPz, Pz; not shown).

